# Bond strength between receptor binding domain of spike protein and human angiotensin converting enzyme-2 using machine learning

**DOI:** 10.1101/2024.04.16.589808

**Authors:** Abdulmateen Adebiyi, Puja Adhikari, Praveen Rao, Wai-Yim Ching

## Abstract

The spike protein (S-protein) of SARS-CoV-2 plays an important role in binding, fusion, and host entry. In this study, we have predicted interatomic bond strength between receptor binding domain (RBD) and angiotensin converting enzyme-2 (ACE2) using machine learning (ML), that matches with expensive *ab initio* calculation result. We collected bond order result from *ab initio* calculations. We selected a total of 18 variables such as bond type, bond length, elements and their coordinates, and others, to train ML models. We then trained five well-known regression models, namely, Decision Tree regression, KNN Regression, XGBoost, Lasso Regression, and Ridge Regression. We tested these models on two different datasets, namely, Wild type (WT) and Omicron variant (OV). In the first setting, we used 90% of each dataset for training and 10% for testing to predict the bond order. XGBoost model outperformed all the other models in the prediction of the WT dataset. It achieved an R2 Score of 0.997. XGBoost also outperformed all the other models with an R2 score of 0.9998 in the prediction of the OV dataset. In the second setting, we trained all the models on the WT (or OV) dataset and predicted the bond order on the OV (or WT) dataset. Interestingly, Decision Tree outperformed all the other models in both cases. It achieved an R2 score of 0.997.

## 1. Introduction

The COVID-19 pandemic started in November 2019, taking millions of lives globally. The severe acute respiratory syndrome coronavirus-2 (SARS-CoV-2) has several variants of concerns (VOC) such as Alpha [1], Beta [2], Delta [3], Gamma [4], and Omicron [5] and variants of interest (VOI) such as Eta [6], Iota [7], Kappa [8], Lambda [9], and Mu [10]. These VOC and VOI have shown the nature of rapidly mutating SARS-CoV-2. With overwhelming effort of the scientific community, the development of vaccines has saved billions of lives. In addition to ongoing research in medicine and biology, scientists from various discipline have collaborated in an effort to enhance prepardness of such a situation in the future.

SARS-CoV-2 is composed of four proteins: spike (S), envelope (E), membrane (M), and nucleocapsid (N) proteins (see **Figure 1(a))**). Among these four proteins, spike protein (S-protein) plays an important role and initiates the infection by binding with human angiotensin converting enzyme 2 (ACE2). S-protein has two subunits S1 and S2. S1 consists of signal peptide (SP), n-terminal domain (NTD), receptor binding domain (RBD), subdomain (SD2). Similary, S2 consists of fusion peptide (FP), heptad repeat 1 (HR1), central helix (CH), connecting domain (CD), heptad repeat 2 (HR2), transmembrane domain (TM), and cytoplasmic tail (CT). Among these domains of S-protein RBD binds with ACE2 (see **Figure 1(b)**).

**Figure 1:**
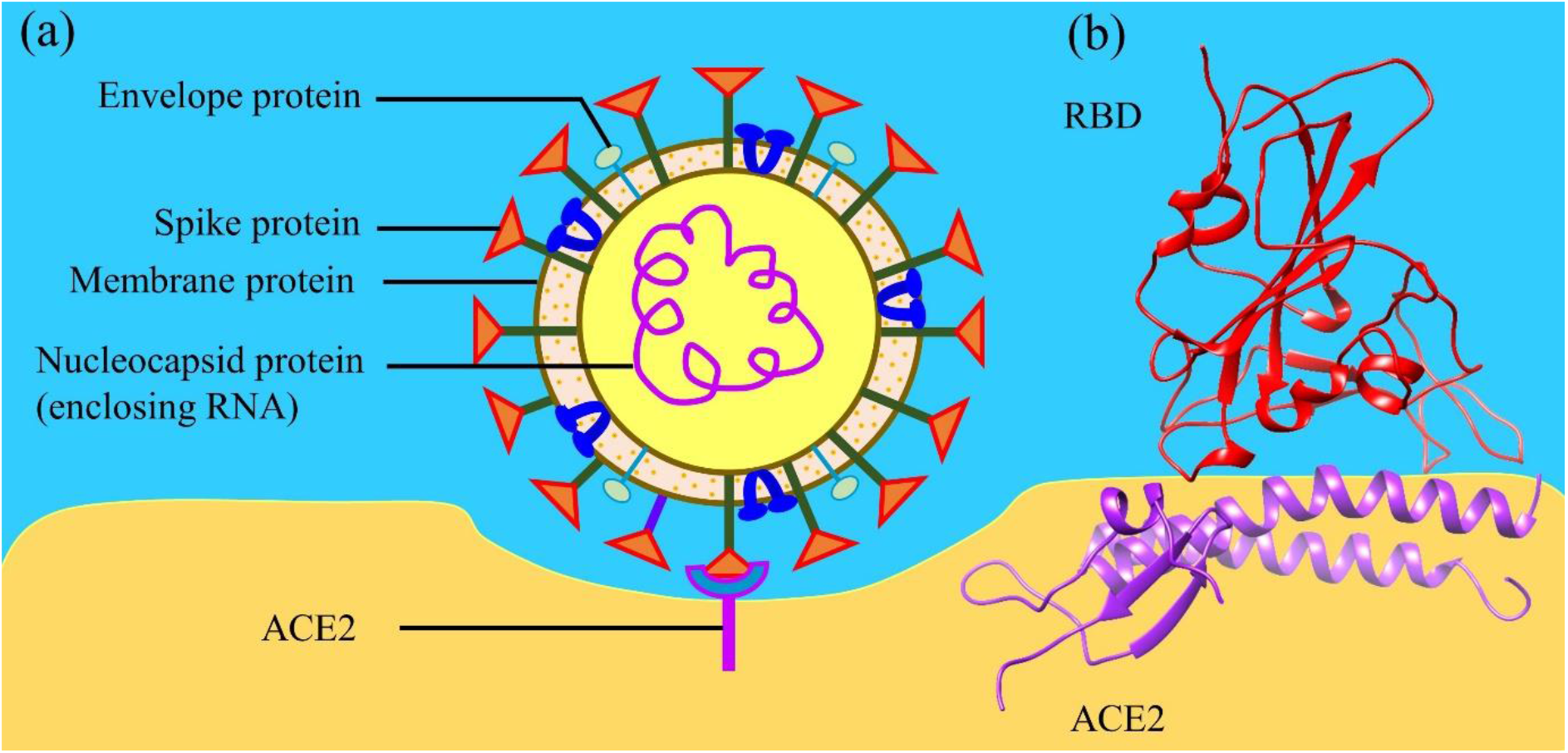
**(a)** Structure of SARS-CoV-2 interacting with angiotensin converting enzyme 2 (ACE2). **(b)** Ribbon structure of interface between receptor binding domain (RBD) and ACE2.

**Figure 2:**
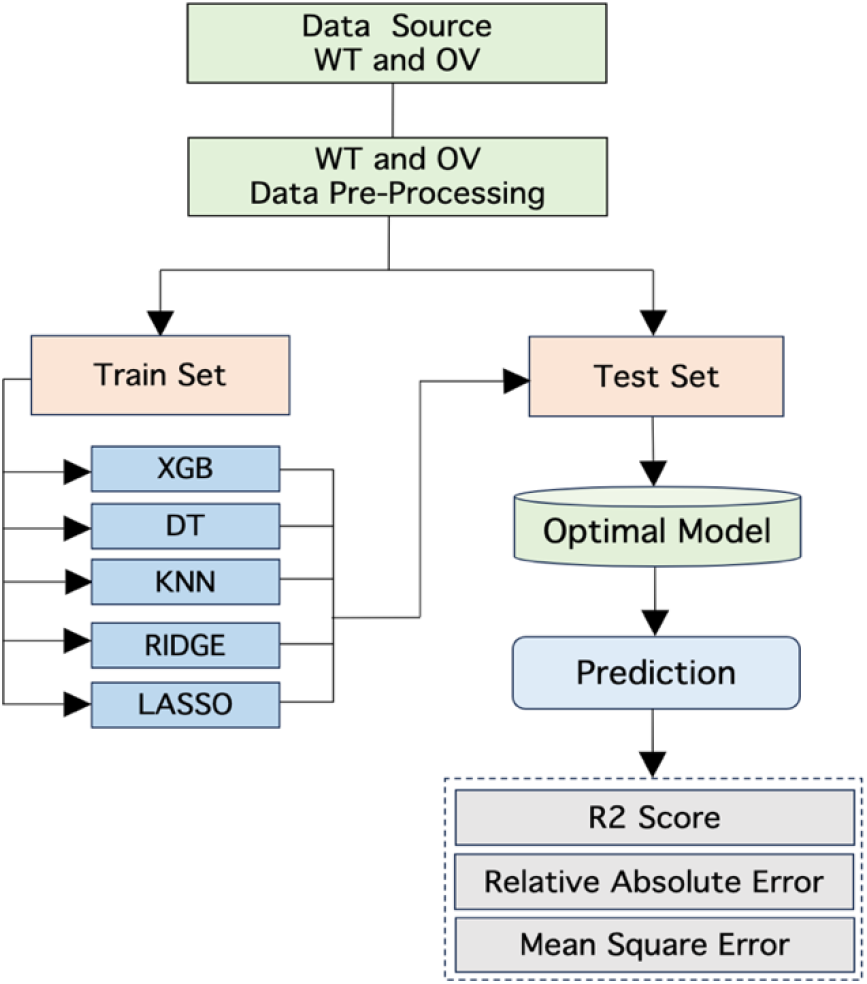
Overall approach for BO prediction

There has been a lot of research including experimental [11-13] and computational [14-18] study focused on RBD-ACE2 interaction. In the computational research, most of the calculations are performed utilizing molecular dynamics (MD) [15, 19-21], with a few employing *ab initio* method. *Ab initio* calculations are known for their accuracy however, they are difficult and expensive especially for such large and complex biomolecules. In our past studies, we have conducted *ab initio* calculations for around 4999 atoms [18], which is very large but not sufficient to calculate biomolecues with several thousands of atoms. Hence, we have utilized *ab initio* methods on such biomolecules using the *divide-and-conquer* strategy. Using this strategy, we divide the complex biomolecules in small sections and calculate their properties such as electronic structure, interatomic bonding, and partial charge. Among these properties interatomic bonding is a crucial one as it reflects the types of bonds involved, and their corresponding strength. Investigating the interatomic bonding within the RBD-ACE2 interface identifies key interacting amino acids. This information can be used to target specific amino acids. Given that RBD-ACE2 is the initial point of contact between the SARS-CoV-2 and human cells, the strategic targeting of these interacting amino acids can disrupt their interaction and stop virus attack. However, *ab initio* calculations can be costly. Therefore, finding ways to achieve comparable accuracy at a reduced expense is essential, and this is where machine learning (ML) steps in.

ML represents a sub-field within artificial intelligence (AI), employing algorithms to discern significant patterns and correlations within complex datasets, facilitating the prediction of specific properties [22]. Using ML, computers can excel in several tasks that humans can perform using their knowledge and wisdom. ML delves into how computer system mimic human learning processes and explores methods for self-improvement to acquire new knowledge [23]. Widely utilized across diverse domains such as healthcare, finance, marketing, and telecommunications, ML models have become indispensable in today’s real-world applications [23].

ML models have also been applied in physics and chemistry. For instance, Hansen *et al*. applied ML techniques to predict molecular properties [24]. The precise and efficient prediction of such properties is crucial for rational computer design in chemical and pharmaceutical industries. Du *et al*. proposed a method for learning the relativistic continuum mass table by using the kernel ridge regression [25]. They used it to learn the nuclear mass table obtained by the relativistic continuum Hartree-Boogoliubov theory. Recently, Adhikari *et al*. suggested the use of ML techniques for predicting potential unknown COVID-19 mutations [26].

In this study, we employed interatomic bonding result obtained through *ab initio* calculations and tested ML to predict these already obtained bonding outcomes. Our objective is to replicate results obtained via *ab initio* methods, with the eventual aim of overcoming the cost limitations associated with such calculations.

## 2. Method

### 2.1 Modelling RBD-ACE2 interface

The interface structures were extracted from the PDB ID 6M0J [13] for the Wild type (WT) and PDB ID 7WBP [27] for the Omicron variant (OV). Amino acids within the sequence S19-I88 and G319-T365 were incorporated from ACE2, while the sequence T333-G526 was included from the RBD. The entire model consists of 311 amino acids. Hydrogen atoms were added using the Leap module with ff14SB force field in the AMBER package [28] resulting into a total of 4817 and 4873 atoms for WT and OV, respectively. The RBD-ACE2 of the OV contains 15 mutations.

### 2.2 Ab initio packages

For the *ab initio* calculations, we have used two density functional theory (DFT) packages—Vienna *ab initio* simulation package (VASP) [29] and orthogonalized linear combination of atomic orbital (OLCAO) [30]. VASP was used for interface model optimization. In VASP, we used projector augmented wave (PAW) [31, 32] method with Perdew-Burke-Ernzerhof (PBE) [31] exchange correlation functional within the generalized gradient approximation (GGA). We used the energy cut off 500 eV with electronic convergence of 10-4 eV, force convergence for ionic relaxation to −10−2 eV, and a single kpoint.

The optimized structure from VASP is used as an input for the OLCAO package. The combination of VASP with OLCAO is very effective for complex biomolecules [14, 16, 18, 30, 33, 34]. OLCAO uses atomic orbitals for basis function expansion. It is used to calculate the interatomic bonding in terms of bond order (BO). BO determines the strength of the bond. OLCAO uses Mulliken’s population analysis to calculate BO. BO is the overlap population *ρ*_*αβ*_between pair of atoms (*α, β*) defined as:

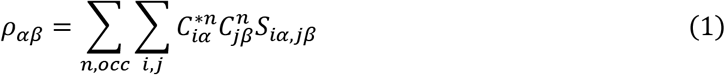

where *S*_*iα,jβ*_ are the overlap integrals between the *i*^*th*^orbital in *α*^*th*^atom and the *j*^*th*^ orbital in *β*^*th*^ atom, and 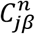 are the eigen vector coefficients of the *n*^*th*^ band, *j*^*th*^orbital in the *β*^*th*^ atom. The dataset for WT and OV obtained from OLCAO were further used as described in section 2.3 below.

### 2.3 Dataset

The BO result obtained from OLCAO calculation served as the source data. The BO prediction was conducted on both interfaces: RBD-ACE2 WT and RBD-ACE2 OV datasets. Note that the RDB-ACE2 WT and RDB-ACE2 OV datasets contained 25,356 rows and 25, 753 rows, respectively. Each dataset had 21 variables. The datasets contained the X, Y and Z coordinates of the bond between the Atom 1 and Atom 2. The variables in the datasets are briefly explained in the table below. We used these datasets to predict the bond strength also known as the BO.

### 2.4 Data Pre-processing

In the original dataset, the total number of variables were 21. We dropped Atom1 and Atom2 as they are both identifiers of the OLCAO package. Eighteen of them were used for training the models, and BO was the target variable for prediction. We used the variables bond, bond length, Ele1, x1, y1, z1, PDBele1, AA1, SeqNo1, ChainN1, Ele2, x2, y2, z2, PDBele2, AA2, SeqNo2 and ChainN2 as input features for our regression models. Label encoding was used to convert the categorical variables to numerical ones before passing them to the ML models. Label encoding is an efficient way to transform categorical data to numerical ones. It is simple, efficient and helps with memory efficiency.

### 2.5 Machine learning

In our work, five well-known ML models were used for the BO prediction. The models were XGBoost Regression, K-Nearest Neighbor (KNN) Regression, Decision Trees Regression, Lasso Regression, and Ridge Regression. They have performed well on different datasets and are commonly used in research projects. XGBoost has outperformed so many models in literature via extreme gradient boosting [35]. Linear regression is used for its simplicity before using other complex models. Decision Trees are often used for many problems because of its interpretability. It can learn complex relationship between features. Next, we briefly describe each model and its unique characteristics for learning on data.

#### Lasso Regression

It is a type of linear regression that combines both linear regression and lasso hyperparameter tuning. It helps to pick the best regression model by discarding less relevant features [36]. The predicted BO is calculated as the product of the input features with the learned coefficients, and then adding the regularization term to penalize the absolute values of the coefficient.

#### Ridge Regression

It is a type of linear regression that uses Ridge Optimization and includes a regularization term to the regression function. The term penalizes the sum of the squared coefficients [37]. The predicted BO is calculated by multiplying the input features with the learned coefficients and then adding the regularization term that penalizes the squared magnitudes of the coefficients.

#### Decision Tree Regression

This regression type uses a tree-like model of decisions to predict the target value. The tree structure has the root node, decision nodes, and leaf nodes. The root node is the initial node of the decision tree. The predicted BO value is computed as the average numerical value for the training data that is stored in the selected leaf node. The optimal structure for the tree is learned during the training process.

#### KNN Regression

It is a supervised learning regression model in which the model predicts a continuous target variable based on the average or weighted average of the values of its k nearest neighbours. The predicted BO is computed by estimating the weighted average of the target values of the k-nearest neighbors.

#### XGBoost

It is a machine learning models that combines decision trees, ensemble learning, and gradient boosting. It is scalable and can be used to train on large datasets. The predicted BO is the sum of individual tree predictions multiplied by the learning rate, which is a hyperparameter used during training. Hyperparameters are the type of parameters that are used to control and manage an ML model during training.

In our experiments, we used the grid search approach to identify the best hyperparameters for the models. We also performed a 10-fold cross validation on our datasets. Cross validation is a technique that is used to evaluate a model by dividing the dataset into two segments [38, 39]. The first segment is used to train the model, and the other segment is used to test the model performance[39]. This process is repeated a number of times so that every data sample is used in the test set leading to a more robust evaluation of the model.

### 2.6 Bond Order Regression

Before the regression process, we first pre-processed our datasets so that it can be easily passed to the ML models. We converted the data that are not in numerical form to numerical form using label encoding. We then split our datasets into the training set and the testing set. We used four different settings in our work:

A. Train on 90% of the RBD-ACE2 WT dataset and evaluate the models on the remaining 10% of the dataset to predict the BO.
B. Train on 90% of the RBD-ACE2 OV dataset and evaluate the models on the remaining 10% of the dataset to predict the BO.
C. Train on the entire RBD-ACE2 WT dataset and evaluate the models on the entire RBD-ACE2 OV dataset to predict the BO.
D. Train on the entire RBD-ACE2 OV dataset and evaluate the models on the entire RBD-ACE2 WT dataset to predict the BO.

## 3. Result

In this section, we present the performance of the aforementioned ML models on the RBD-ACE2 WT and RBD-ACE2 OV datasets. We implemented all the models in Python using Scikit Learn [40], Numpy [41], Matplotlib, and XGBoost libraries [42]. We used the best model after the hyperparameter tuning for each approach for the BO prediction.

### 3.1 Performance Metrics

Next, we will discuss the different performance metrics that were used in this work to evaluate our models. Our regression models aimed to predict the BO in the chosen datasets. We evaluated the models based on the R2 Score, Root Mean Square Error (RMSE), and the Relative Absolute Error metrics (RAE), which are explained below.

#### R2

This metric is referred to as the goodness of fit or the coefficient of determination. It shows how the regression line approximates the actual data. The values are between 0 to 1 with 1 being the value when the model fits the data perfectly.

#### RAE

The RAE is the ratio of the mean error (residuals) to the errors by the naïve model.

#### RMSE

This is calculated by computing the square root of the Mean Squared Error (MSE). MSE is the average of the squared differences between the predicted and expected target values. If all the predicted BO values are the same as the true BO values, RSME will be 0. RMSE is a good metric for regression analysis because it penalizes large error. The RMSE measures the standard deviation of the errors when the regression model makes a prediction.

First, we present the results for Settings (A) and (B). **Table 2** shows the performance metrics of the various ML models for Setting (A). XGBoost outperformed all the other models. It achieved an R2 score of 0.997, RAE of 0.029, and RMSE of 0.008. **Table 3** shows the performance metrics of the various ML models for Setting (B). Once again, XGBoost outperformed other models and achieved an R2 score of 0.998, RAE of 0.028, and RMSE of 0.006.

**Table 1:**
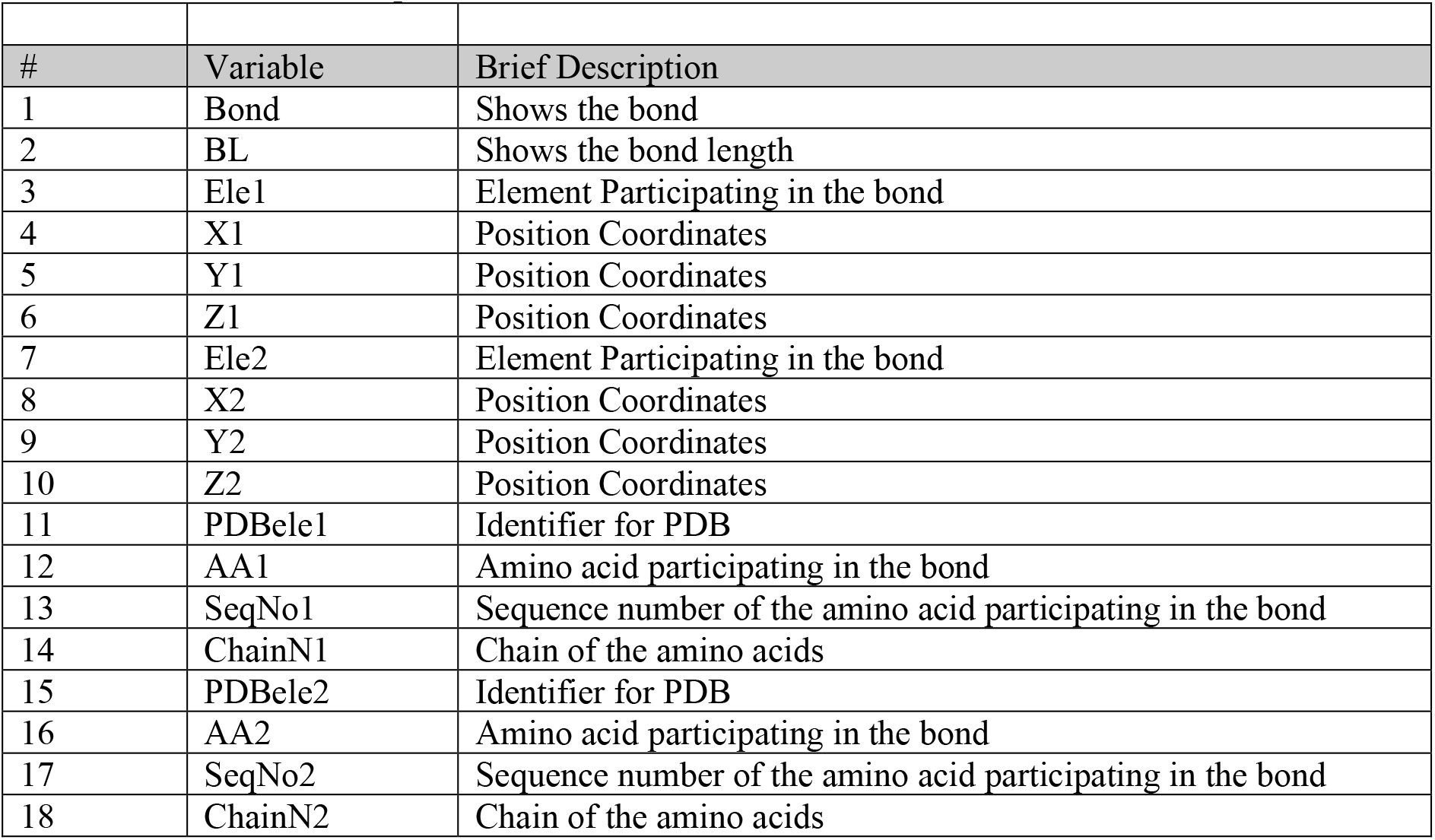
Brief description of the variables.

**Table 2:** Performance Metrics of the ML models for Setting (A) (best value is shown in **bold)**

| Models | R2 | RAE | RMSE |
| --- | --- | --- | --- |
| XGBoost | **0.997** | 0.029 | **0.008** |
| Decision Trees | 0.996 | **0.025** | 0.009 |
| KNN | 0.320 | 0.800 | 0.130 |
| Ridge Regression | 0.830 | 0.392 | 0.065 |
| Lasso Regression | 0.442 | 0.696 | 0.118 |

**Table 3:** Performance Metrics of the ML models for Setting (B) (best value is shown in **bold)**

| Models | R2 | RAE | RMSE |
| --- | --- | --- | --- |
| XGBoost | **0.998** | 0.028 | **0.006** |
| Decision Trees | 0.996 | **0.025** | 0.009 |
| KNN | 0.339 | 0.726 | 0.126 |
| Ridge Regression | 0.826 | 0.405 | 0.065 |
| Lasso Regression | 0.444 | 0.712 | 0.116 |

Next, we present the results for Settings (C) and (D). **Table 4** and **Table 5** show the performance results of our various ML models for Setting (C) and Setting (D), respectively. In both cases, Decision Trees outperformed other models.

**Table 4:** Performance Metrics of the ML models for Setting (C) (best value is shown in **bold)**

| Models | R2 | RAE | RMSE |
| --- | --- | --- | --- |
| XGBoost | 0.995 | 0.049 | 0.011 |
| Decision Tress | **0.997** | **0.028** | **0.008** |
| KNN | 0.386 | 0.708 | 0.121 |
| Ridge Regression | 0.824 | 0.406 | 0.065 |
| Lasso Regression | 0.386 | 0.710 | 0.115 |

**Table 5:**
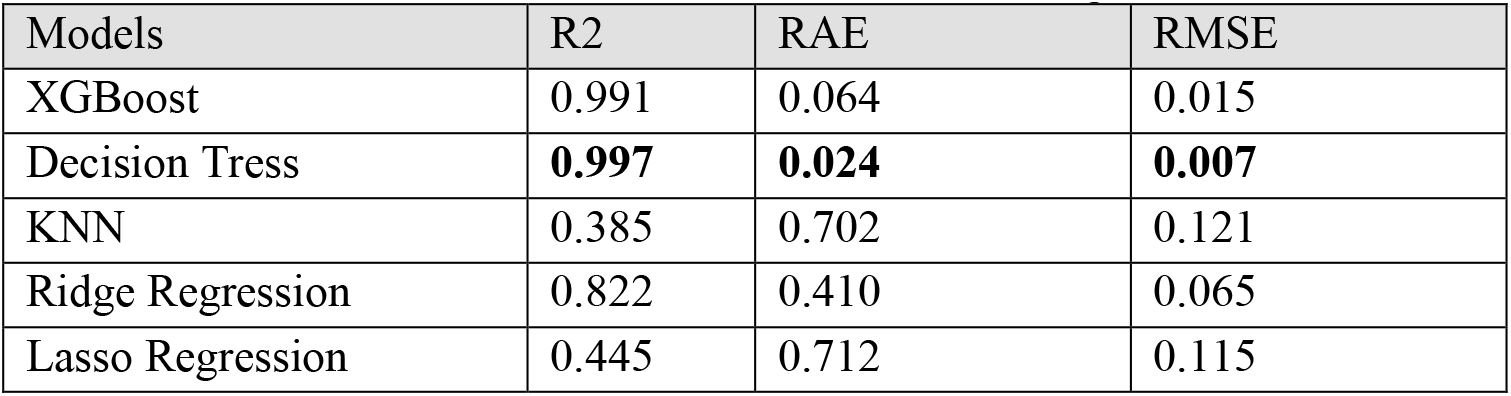
Performance Metrics of the ML models for Setting (D) (best value is shown in **bold)**

| Models | R2 | RAE | RMSE |
| --- | --- | --- | --- |
| XGBoost | 0.991 | 0.064 | 0.015 |
| Decision Tress | **0.997** | **0.024** | **0.007** |
| KNN | 0.385 | 0.702 | 0.121 |
| Ridge Regression | 0.822 | 0.410 | 0.065 |
| Lasso Regression | 0.445 | 0.712 | 0.115 |

We additionally plotted the R2 values for the various models in Settings (A) and (B). These are illustrated in **Figures 3 and 4**, where the x-axis represents the actual BO and the y-axis denotes the predicted BO. For models that achieved high R2 score, we observed that the predictions were clustered around the 45-degree line (shown as a dotted line). Decision Trees and XGBoost models performed very well. For models with low R2 score, we observed that the predicted values were scattered differently. Clearly, the linear regression models were unable to capture the underlying distribution of the BO values. KNN regression also performed poorly on the tested datasets. Similar trends were observed for Settings (C) and (D) where Decision Trees and XGBoost achieved high R2 scores. In the interest of space, the plots are not shown in the paper.

**Figure 3:**
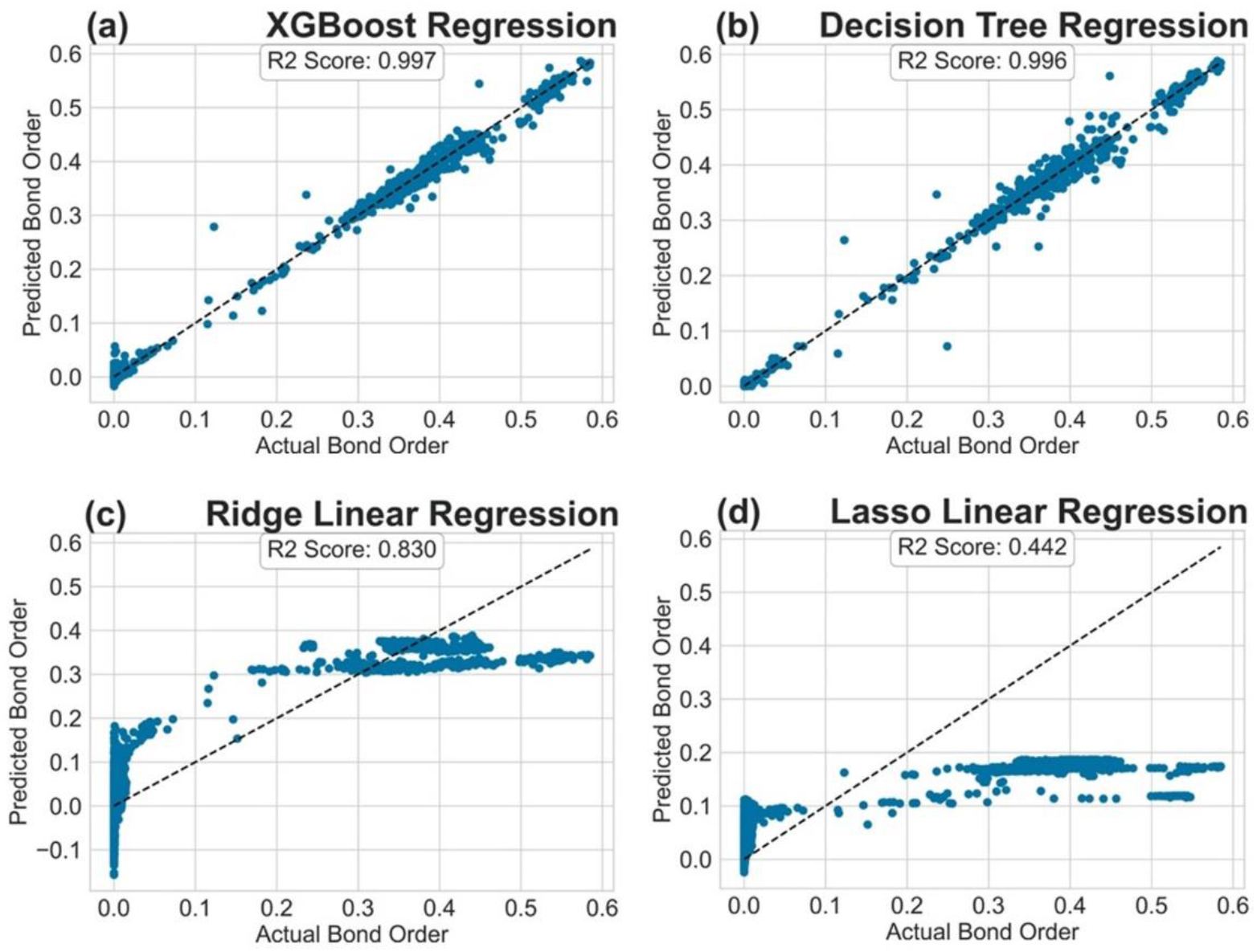
R2 plot for the tested regression models in Setting (A).

**Figure 4:**
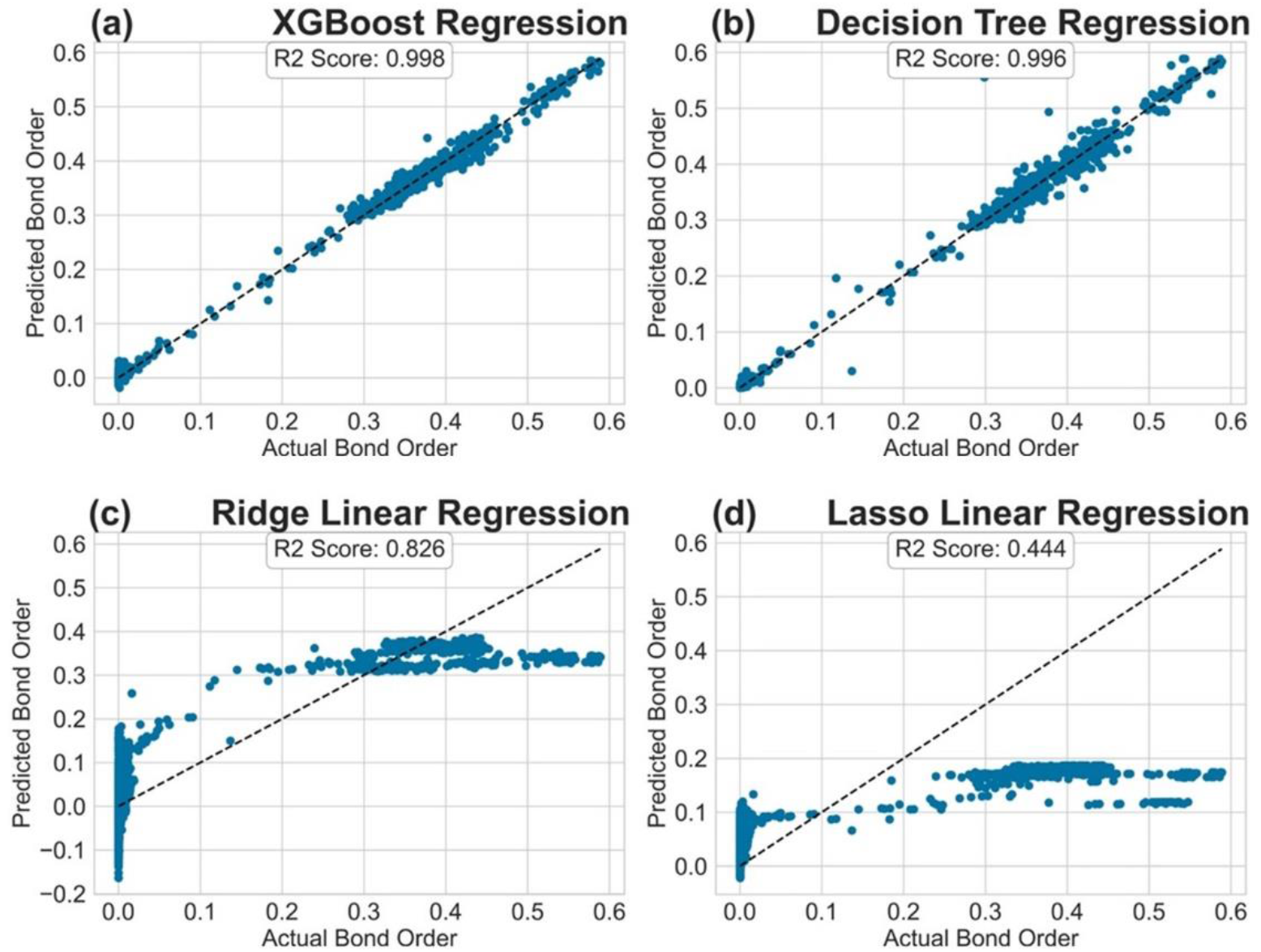
R2 Plot for the tested regression models in Setting (B).

## 4. Conclusion

In this work, we explored the efficacy of well-known ML models in predicting BO values derived from ab initio calculations, which are computationally demanding to simulate on high performance computing platforms. Specifically, we employed XGBoost Regression, Decision Tree Regression, KNN Regression, Lasso Regression, and Ridge Regression for BO prediction. The XGBoost model outperformed all other models when the training and testing samples were from the same dataset (i.e., Settings (A) and (B)). However, Decision Trees outperformed all other models when the training and testing samples were from different datasets (i.e., Settings (C) and (D)). Linear regression models and KNN regression performed poorly in all settings. Our investigation shows that ML models such as Decision Trees and XGBoost can be used to accurately predict the BO values without expensive simulation-based computations.

## Acknowledgements

This research used the resources of the National Energy Research Scientific Computing Center (NERSC), a DOE office of Science User Facility supported by the Office of Science of the U.S. Department of Energy by U.S. Department of Energy under the contract number DE-AC03-76SF00098, DE-AC02-05CH11231 using NERSC award NERSC DDR-ERCAP0023727, and the Research Computing Support Services (RCSS) of the University of Missouri System. The third author (P. R.) would like to acknowledge the support of the National Science Foundation (NSF) Grant No. 2201583.

## Notes

### Competing Interest Statement

The authors have declared no competing interest.

